# Type III secretion system of *Pseudomonas aeruginosa* affects mucin gene expression via NF-κB and AKT signaling in human carcinoma epithelial cells and a pneumonia mouse model

**DOI:** 10.1101/442061

**Authors:** Ji-Won Park, In-Sik Shin, Sei-Ryang Oh, Un-Hwan Ha, Kyung-Seop Ahn

**Author notes:** Author to whom correspondence should be addressed: Natural Medicine Research Center, Korea Research Institute of Bioscience and Biotechnology, 30 Yeongudanji-ro, Ochang-eup, Cheongwon-gu, Cheongju-si, Chungbuk 28116, Republic of Korea.

## Abstract

The type III secretion system (T3SS) in *Pseudomonas aeruginosa* has been linked to severe disease and poor clinical outcomes in animal and human studies. Of the various T3SS effector genes, ExoS and ExoT showed mutually exclusive distributions, and these two genes showed varied virulence. We aimed to investigate whether the ExoS and ExoT effector proteins of *P. aeruginosa* affect the expression of the proinflammatory mediators Muc7, Muc13, Muc15, and Muc19 via the NF-κB and AKT signaling pathways. To understand the role of the T3SS, we used AExoS, AExoT, and T3SS transcriptional activator ExsA mutants (ExsA∷Ω), as well as A549 cells stimulated with *P. aeruginosa* strain K (PAK). We investigated the effects of ΔExoS, ΔExoT, and ExsA∷Ω on the development of pneumonia in a mouse model and on Muc7, Muc13, Muc15, and Muc19 production in A549 cells. ΔExoS and ΔExoT markedly decreased the neutrophil count in the bronchoalveolar lavage fluid, with a reduction in Muc7, Muc13, Muc15, and Muc19 expression ΔExoS andΔExoT reduced NF-κB and AKT phosphorylation, together with Muc7, Muc13, Muc15, and Muc19 expression in PAK-infected mice and A549 cells. In conclusion, *P. aeruginosa* infection induced the expression of Mucus, and the *P. aeruginosa* T3SS appeared to be a key player in Muc7, Muc13, Muc15, and Muc19 expression, which is further controlled by NF-κB and AKT signaling. These findings might be useful to devise a novel therapeutic approach for the treatment of chronic pulmonary infections by targeting ExoS and ExoT.

**Author Summary:** *Pseudomonas aeruginosa* is a ubiquitous gram-negative bacterium causing serious infections. Many clinical isolates of *P. aeruginosa* have a specialized apparatus for injecting toxins into eukaryotic cells, namely, the type III secretion system (T3SS). The T3SS is a syringe-like apparatus on the bacterial surface, with 4 effector toxins: ExoS, ExoT, ExoY, and ExoU. We investigated the effect of ExoS and ExoT of the T3SS of *P. aeruginosa* K strain (PAK). Mucus plays a vital role in protecting the lungs from environmental factors, but conversely, in muco-obstructive airway disease, mucus becomes pathologic. We showed that infection with ExoS and ExoT induced Muc7, Muc13, Muc15, and Muc19 expression in host cells. PAK clinical strains induce proinflammatory cytokine production through the T3SS, and this involves NF-κB and SP1/AKT activation in pneumonia mouse models. Mucus induction in response to ExoS and ExoT infection relied on NF-κB and SP1/AKT activation. Our findings highlight the roles of Muc7, Muc13, Muc15, and Muc19 in inducing proinflammatory cytokine expression during ExoS and ExoT exposure in PAK infections, paving the way for a novel therapeutic approach for the treatment of pulmonary infections.

## Introduction

The gram-negative bacterium *Pseudomonas aeruginosa* uses a complex type III secretion system (T3SS) to inject effector proteins into host cells [1,2]. The T3SS is a major virulence determinant that manipulates eukaryotic host cell responses that is present in a broad range of pathogens. It is a specialized needle-like structure that delivers effector toxins directly from the bacterium into the host cytosol in a highly regulated manner [3]. This system is activated on contact with eukaryotic cell membranes, interferes with signal transduction, and causes cell death or alterations in host immune responses [4]. The T3SS in *P. aeruginosa* has been linked to severe disease and poor clinical outcomes in animal and human studies [4,5]. The features of this interesting secretion system have important implications for the pathogenesis of *P. aeruginosa* infections and for other T3SSs. *P. aeruginosa* has four known effector toxins: ExoS, ExoT, ExoY, and ExoU. These proteins can modify signal transduction pathways and counteract innate immunity [6,7]

Mucins are a major component of the respiratory mucus. Mucins are either membrane-bound (like MUC1) with a role in sensing external information and transducing it to cells, or secreted, the type of which is characterized by a high molecular weight and viscosity. They are glycoproteins secreted by the mucosal and submucosal glands. The mucin molecule consists of a polypeptide core with branched oligosaccharide side chains, each of which contains 8 to 10 sugars [8,9]. Molecular cross-linking of this structure contributes to the viscoelastic property of mucus [10]. Despite their recalcitrance, mucins are a main nutrient source for niche-specific microbiota of the gut and oral cavity. For example, oral streptococci produce a variety of glycolytic and proteolytic enzymes that liberate bioavailable carbohydrates from salivary glycoproteins [11,12]. At least 20 human mucin genes have been identified by cDNA cloning: *MUC1*, *MUC2*, *MUC3A*, *MUC3B*, *MUC4*, *MUC5AC*, *MUC5B*, *MUC6*, *MUC7*, *MUC8*, *MUC9*, *MUC12*, *MUC13*, *MUC15*, *MUC16*, *MUC17*, *MUC19*, *MUC20*, *MUC21*, and *MUC22*) [13]. However, these mucins cannot maintain homeostasis in intrabronchial respiratory epithelial cells of patients with weak immunity, and when overexpressed, cause mucous membrane damage due to excessive secretion of mucus, and need to be blocked. However, the underlying mechanisms have been clearly clarified for only a few mucins to date. Muc7, Muc13, Muc15, and Muc19 have been implicated in bronchial inflammation among mucin targets, but the mechanism of the mechanism has not been studied to date. Especially in immunocompromised patients and children infected with the opportunistic pathogen *P. aeruginosa*, the risk of secondary infection is high and leads to not only asthma and airway hypersensitivity as well as deterioration of existing disease, but also mucus accumulation, which can result in terrible death due to difficulty of breathing by chronic obstructive pulmonary disease [14,15]. However, the mechanism underlying this condition is yet to be clarified.

NF-κB is also a high expression level transcription factor involved in many inflammation’s formation and development [16]. Moreover, SP1/NF-κB pathway has been reported connected to cell migration, invasion and EMT (epithelial to mesenchymal transition) recently [17,18]. Furthermore, SP1 and NF-κB (p65) were found significantly upregulated in ExoS and ExoT infected cell. The ExsA::Ω and ΔST does not increased the expression level of SP1 and NF-κB (p65) in vitro and in vivo. The expression level of Muc7, Muc13, Muc15, and Muc19 may be increased through activation of SP1 and NF-κB (p65) due to infection of ExoS and ExoT.

Here, we investigated the effect of ExoS and ExoT of *P. aeruginosa* strain K (PAK) on the induction of Muc7, Muc13, Muc15, and Muc19 expression, as well as the underlying mechanism, in host cells and a pneumonia mouse model. We expected our study to provide new insights into the roles of Muc7, Muc13, Muc15, and Muc19 in inducing proinflammatory cytokine expression in response to ExoS and ExoT exposure in PAK infection.

## Materials and Methods

### Bacterial strains

All strains and plasmids used in this study are listed in Supporting information S1 Table. Chromosomal mutants were all derived from the same parental PAK strain, as indicated in S1 Table, and were generated by allelic exchange (for details, see the Supporting information S1 Text). PAK strains vary widely in their expression of virulence genes. The strain used in this study strongly expresses the T3SS genes. Regions flanking the appropriate mutation were amplified using chromosomal DNA as a template (unless specified otherwise), joined by “splicing by overlap extension PCR,” and cloned into the appropriate plasmid using the indicated restriction enzymes. PAK-ΔST is a *P. aeruginosa* clinical isolate that naturally carries the ExoY gene, but lacks the genes for ExoS and ExoT. We reported that the pUCP18-PAKexoS (S) and pUCP18-PAKexoT (T) mutant strains are secretion competent and export the expected effector proteins [19]. Antibiotics were used when necessary at the following concentrations: for plasmids in *Escherichia coli*, 50 μg/mL ampicillin, 15 μg/mL gentamicin, and 25 μg/mL kanamycin; for *P. aeruginosa*, 500 μg/mL carbenicillin, 100 μg/mL gentamicin, and 100 μg/mL tetracycline.

### Human cell culture

A549 adenocarcinomic human alveolar basal epithelial cells, and H292 human airway epithelial cells were purchased from the American Type Culture Collection (ATCC, Manassas, VA, USA). The cells were maintained in RPMI1640 (Invitrogen, Grand Island, NY, USA) supplemented with 10% fetal bovine serum (FBS; Invitrogen) in the presence of penicillin (100 U/mL), streptomycin (100 μg/mL; Sigma-Aldrich, St. Louis, MO, USA), and HEPES (25 mM), at 37 °C in a 5% CO_2_ atmosphere.

### *In-vitro* bacterial infection

For direct bacterial challenge of A549 cells, bacterial strains were grown in tryptic soy broth (Sigma-Aldrich) at 37 ° C until the OD_600_ reached 1. The bacterial cultures were centrifuged at 7,000 × *g* for 10 min, washed with PBS, and resuspended at a ratio of 1:20 bacterial cells to A549 cells or H292 cells.

### qRT-PCR analysis

Total RNA was extracted using TRIzol^®^ reagent (Invitrogen) following the manufacturer’s protocol and was used to synthesize cDNA using the AMPIGENE^®^ cDNA Synthesis Kit (Enzo Life Sciences, NY, USA). PCRs were conducted using SYBR Green PCR Master Mix (KAPA Biosystems, Woburn, MA, USA) and the primers listed in S2 Table. Reactions were run in a CFX96 real-time PCR system (Bio-Rad, Hercules, CA, USA) using the following thermal conditions: stage 1, 50°C for 2 min and 95°C for 10 min; stage 2, 95°C for 15 s and 60°C for 1 min. stage 2 was repeated for 40 cycles. Relative mRNA levels were calculated using the comparative CT method and normalized to the level of GAPDH.

### Immunoblot analysis

Cells were lysed with 20 mM Tris-HCl (pH 7.4), 50 mM NaCl, 50 mM sodium phosphate, 30 mM NaF, 5 μM zinc chloride, 2 mM iodoacetic acid, and 1% Triton X-100 for 10 min at room temperature on ice for 20 min with regular vortexing. The lysate was centrifuged at 16,000 × *g* for 10 min at 4 ° C and the supernatant was collected. The protein concentration in the supernatant was measured using bicinchoninic acid (Pierce, Rockford, IL, USA). Proteins were separated by SDS-PAGE and transferred to polyvinylidene difluoride membranes.

Membranes were blocked in TBS (10 mM Tris-HCl (pH 7.5), 150 mM NaCl) containing 5% nonfat dry milk for 2 h and incubated with the primary antibodies anti-Muc7 (Enzo Life Sciences, Farmingdale, NY, USA), anti-Muc13 (Thermo-Fisher Scientific, Waltham, MA, USA), anti-Muc15 (Enzo Life Sciences, Farmingdale, NY, USA), anti-Muc19 (Enzo Life Sciences), anti-p65 (Santa Cruz, Dallas, TX, USA), anti-phospho-p65 (Cell Signaling, Danvers, MA, USA), anti-IκBα (Cell Signaling), anti-phospho-IκBα (Cell Signaling), anti-AKT (Cell Signaling,), anti-phospho AKT (Cell Signaling), anti-SP1 (Santa Cruz) or β-actin (Thermo-Fisher Scientific) for 18 h at 4 °C. The immunoblots were washed and incubated with appropriate secondary antibodies and visualized using SuperSignal™ West Pico Chemiluminescent Substrate (Pierce, Rockford, IL, USA) or SuperSignal™ West Femto Maximum Sensitivity Substrate (Pierce).

### Promoter analysis

SP1 promoter activity was assessed using a luciferase assay system (Promega, Madison, WI, USA) according to the manufacturer’s instructions, and β-galactosidase expression (pCH110) was used for normalization. A549 cells were transfected with pGL4.14 and the indicated luciferase SP1 promoter construct.

A549 cells were transfected with pGL4.43 (luc2P/NF-κB-RE/Hygro) plasmid using Lipofectamine 2000 transfection reagent (Invitrogen/Thermo-Fisher Scientific, Carlsbad, CA, USA) according to the manufacturer’s protocol. Twenty hours after transfection, the cells were stimulated with PAK for 2 h, harvested, and assessed for luciferase activity using the ONE-Glo™ luciferase reporter assay system (Promega) according to the manufacturer’s instructions.

### Immunohistochemistry

H292 cells were cultured on Permanox plastic chamber slides (Nunc, Rochester, NY, USA) and fixed in methanol at 4°C for 20 min. The slides were washed three times with PBS, and blocked with 3% (w/v) BSA in PBS for 30 min. Then, the slides were incubated with anti-NF-κB p65 subunit (rabbit polyclonal IgG, 1:200 dilution, Santa Cruz) antibody at 4°C for 24 h. The slides were washed to remove excess primary antibody and then incubated with anti rabbit Alexa Fluor 488-conjugated secondary antibody (Invitrogen, Carlsbad, CA) for 2 h at room temperature, washed with PBS, and then mounted using ProLong Gold Antifade reagent containing 4’,6-diamidino-2-phenylindole (DAPI) (Invitrogen) for 5 min prior to visualization by confocal laser scanning microscopy (LSM510; Carl Zeiss, Oberkochen, Germany). All samples were photographed under the same exposure conditions, and nuclei were quantified from the images obtained.

### Animal infection

Four-week-old, specific pathogen-free female C57BL/6 mice were purchased from the Orient Co. (Seoul, Korea) and were used after a week of quarantine and acclimatization. The mice were allowed access to sterilized tap water and standard rodent chow. Bacteria were centrifuged and resuspended to the appropriate CFU/mL in PBS as determined by optical density, and plated out in a serial dilution on nutrient broth agar plates. LPS was injected intranasally by dissolving 5 μg in 50 μl of PBS. Mice were slightly anaesthetized by intraperitoneal injection of pentobarbital (Virbac, Fort Worth, TX, USA). Bacterial solutions at appropriate concentration (2.5 × 10^6^ CFU per mouse in 50 μL PBS) were then administered by intranasal instillation (25 μL per nostril). Control mice were inoculated intranasally with 50 μL of PBS. Survival experiments and analysis of bronchoalveolar lavage fluid (BALF) were performed as previously described [20,21]

### Histology

After BALF samples were obtained, the mice were sacrificed by intraperitoneal injection of pentobarbital (50 mg/kg; Hanlim Pharm. Co., Seoul, Korea), and lung tissues were collected and fixed in 10% (v/v) neutral-buffered formalin. The tissues were embedded in paraffin, sectioned at 4-μm thickness, and stained with hematoxylin and eosin solution (hematoxylin, Sigma MHS-16; eosin, Sigma HT110-1-32). Quantitative analysis of inflammation and mucus production was performed in at least 4 squares per slide using an image analyzer (Molecular Devices Inc., Sunnyvale, CA, USA).

### siRNA-mediated knockdown

siRNAs targeting IKKα and IKKβ were designed and synthesized by Dharmacon (Lafayette, CO, USA). A549 cells were resuspended in serum-free DMEM, and 2.5 × 10^5^ cells/mL were seeded in a 12-well plate and transfected with siRNA-IKKα/siRNA-IKKβ or siRNA-negative control (NC), using Lipofectamine RNAiMAX (Invitrogen; Thermo Fisher Scientific, Inc.). After 6 h, the medium was changed with RPMI1640 containing 10% FBS. Cells were harvested at 48 h after transfection for RT-qPCR or at 72 h for western blot analysis. Three different IKKα/IKKβ-specific siRNAs were screened, and the most efficient one was selected for experiments.

### Ethics Statement

All experimental procedures were carried out in accordance with the NIH Guidelines for the Care and Use of Laboratory Animals and were approved by the Korea Institute of Bioscience and Biotechnology Animal Care and Use Committee (IACUC KRIBB-AEC-14094).

### Statistical analysis

Data represent the mean ± standard error of the mean (SD). Statistical differences among groups were determined by one-way ANOVA with repeated measures followed by Newman–Keuls testing in SPSS 14.0 (IBM Software, Armonk, NY, USA). *P* < 0.05 was considered statistically significant.

## Results

### Muc7, Muc13, Muc15, and Muc19 expression is increased in PAK-infected A549 and H292 cells

*P. aeruginosa* PAK affected the expression of Muc7, Muc13, Muc15, and Muc19 in a MOI-and time-dependent manner. As the multiplicity of infection (MOI) of PAK increased, the expression of Muc7, Muc13, Muc15, and Muc19 in A549 cells increased at the mRNA (Fig 1A–D) and the protein (Fig 1I) level. The mRNA expression of all 4 mucins tended to decrease after 12 h, while at the protein level, expression decreased after 4 h (Fig 1E–H, J). Immunohistochemistry (IHC) of H292 cells infected with PAK at MOI 20 confirmed the increase in Muc7, Muc13, Muc15, and Muc19 (Fig 1K). Thus, Muc7, Muc13, Muc15, and Muc19 in A549 cells increased in a concentration- and time-dependent manner by PAK infection.

**Fig 1.**
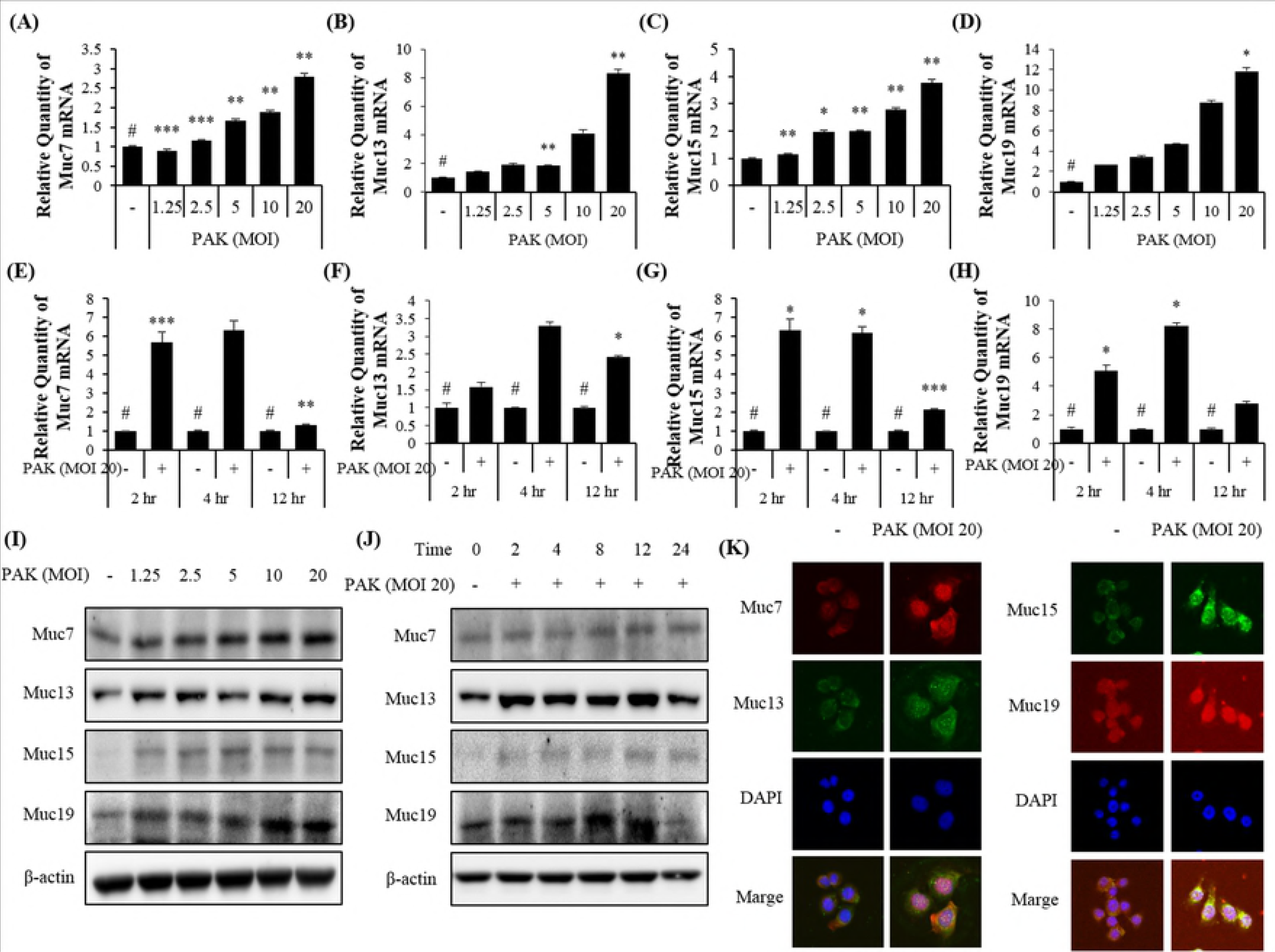
Muc7, Muc13, Muc15, and Muc19 expression in PAK-infected A549 and H292 cells. (A–D) PAK MOI-and (E–H) time-dependent increases in Muc7, Muc13, Muc15, and Muc19 mRNA expression as measured by qRT-PCR. Data are presented as the mean ± SD. “–”, A549 cells treated with PBS; PAK, A549 cells infected with *P. aeruginosa.* #Significantly different from the normal control group, \**P* < 0.05, \*\**P* < 0.01, \*\*\**P* < 0.001 vs. negative control (PBS) Western blots showing PAK MOI- (I) and time-dependent (J) regulation of Muc7, Muc13, Muc15, and Muc19 protein expression. (K) Intracellular staining of Muc7, Muc13, Muc15, and Muc19 in PAK-infected H292 cells (magnification, 400×). Nuclei were stained with DAPI.

### Effects of NF-κB signaling inhibitors BAY11-7082 and LY-294002 on Muc7, Muc13, Muc15, and Muc19 expression and NF-κB p65 and AKT phosphorylation in PAK-infected A549 cells

Bay11-7082 is an IκBα inhibitor and LY-294002 is a well-known inhibitor of PI3K signaling. We previously reported that PAK increased p65 phosphorylation in an epithelial cell line [22]. In addition, PAK reportedly increased AKT phosphorylation lung cancer cells [23,24]. Therefore, we tested the effects of the above inhibitors on mucin expression in A549 cells infected by PAK. Bay11-7082 and LY-294002 suppressed the increases in Muc7, Muc13, Muc15, and Muc19 mRNA expression induced by PAK (Fig 2A–D). The effect of PAK on Muc7 expression was hardly affected by the inhibitors while Muc13 expression was less affected. Muc15 and Muc19 were more strongly suppressed in the presence of LY-294002 than in the presence of Bay11-7082. Similar findings were obtained for protein expression by western blotting (Fig 2E).

**Fig 2.**
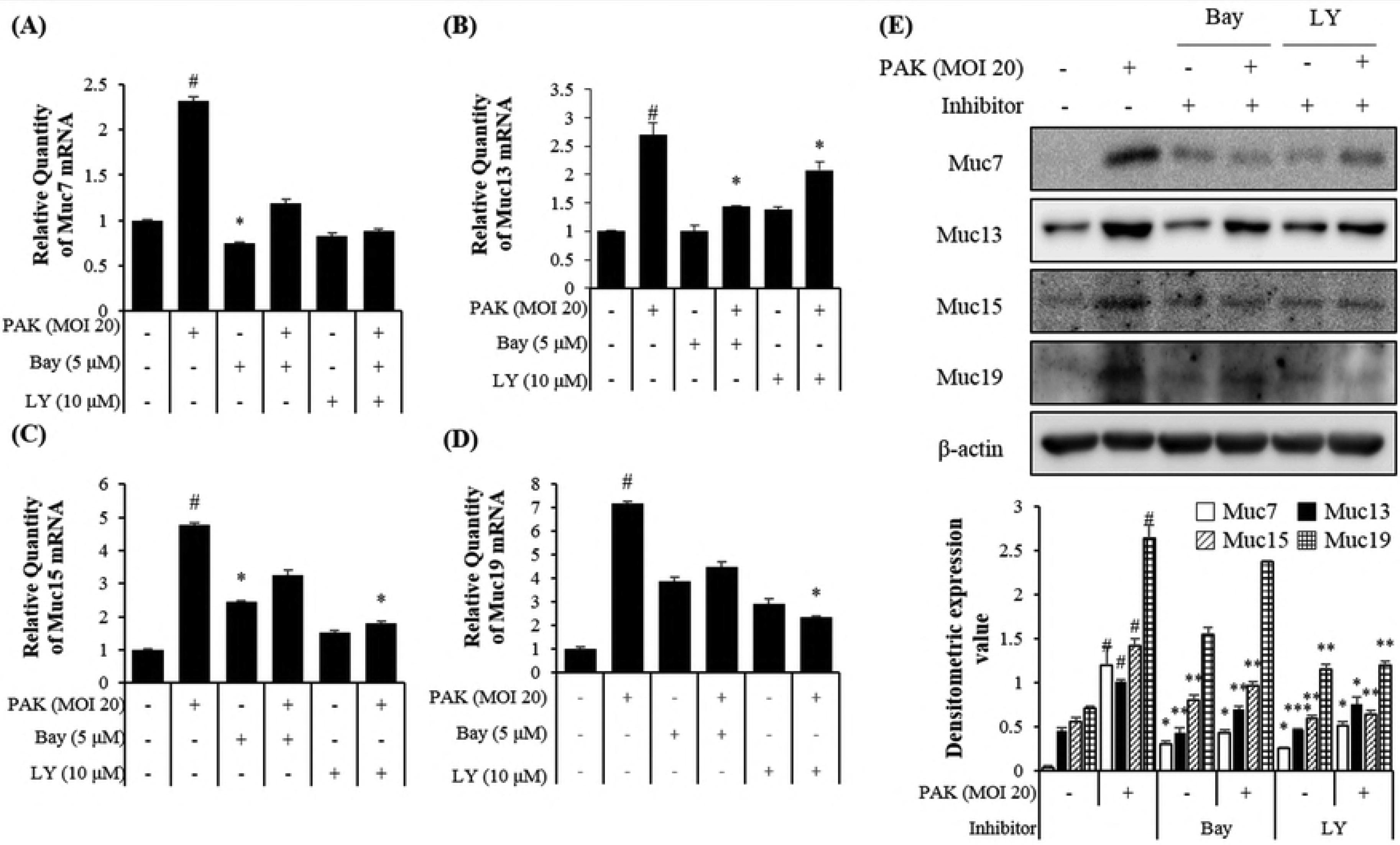
Inhibitory effects of BAY-11-7082 and LY-294002 on the expression of Muc7, Muc13, Muc15, and Muc19 in PAK-infected A549 cells. A549 cells were pretreated with Bay11-7082 or LY-294002 and then infected with PAK (MOI 20) for 4 h. (A–D) Muc7, Muc13, Muc15, and Muc19 mRNA expression as assessed by qRT-PCR. (E) Western blot of Muc7, Muc13, Muc15, and Muc19 protein expression and quantitative data. Western blot bands were quantified using ImageJ software. The data are presented as the mean ± SD. “–”, A549 cells treated with PBS; PAK, A549 cells infected with *P. aeruginosa*; Bay, Bay11-7082; LY, LY-294002; #Significantly different from the normal control group, \**P* < 0.05; \*\**P* < 0.01; \*\*\**P* < 0.001, significantly different from the respective controls.

Phosphorylation of NF-κB and AKT was assessed in PAK-infected A549 cells in the presence or absence of each inhibitor. In PAK-infected A549 cells, luciferase assay indicated that transcription factor activities of NF-κB and SP1 increased. In the presence of Bay11-7082 and LY-294002 these PAK-induced transcription factor activities of NF-κB and SP1 were significantly suppressed as indicated by luciferase analysis and western blot analysis (S1A–D Fig). In PAK-infected A549 cells, the proinflammatory cytokines IL-8 and IL-6 increased, while Bay11-7082 and/or LY294002 significantly suppressed this response as indicated by qRT-PCR and ELISA of IL-6 and IL-8 (S2A–D Fig). Thus, PAK-induced NF-κB and SP1 signal transduction as well as IL-8 and IL-6 of proinflammatory cytokine expression is suppressed by the use of the inhibitors BAY11-7082 and LY-294002 in A549 cells.

### ExsA::Ω mutant does not induce expression of Muc7, Muc13, Muc15, and Muc19 via NF-κB p65 and AKT phosphorylation in PAK-infected A549 cells

ExsA::Ω is a mutant that has no functional T3SS. We compared the induction of expression of the 4 mucins in A549 cells between PAK and exsA::Ω. As expected, mRNA expression of all 4 mucins was significantly induced by wild-type PAK, while exsA::Ω did not induce their expression as compared to non-infected control cells (Fig 3A–D). Similar findings were obtained for mucin protein expression (Fig 3A–D). AKT and SP1 phosphorylation was less strongly induced by exsA::Ω than by wild-type PAC as indicated by western blotting (Fig 3E). SP1 and NF-κB transcription factor activities were significantly induced by wild-type PAC but not by exsA::Ω as indicated by luciferase assays (Fig 3F, H). Similar to AKT and SP1 phosphorylation, p65 and IκBa phosphorylation was less strongly induced by exsA::Ω than by wild-type PAC as indicated by western blotting (Fig 3I). IHC confirmed the findings for SP1 and p65 (Fig 3G, J). There are multiple reports that PAK affects NF-κB signaling, but there was no report that PAK phosphorylated p65, IκBa and SP1 to affect Muc7, Muc13, Muc15, and Muc19 in experiments using A549 cells. Taken together, these results indicated that a functional T3SS is required for PAK to exert its effects on SP1/AKT signaling, NF-κB signaling, and mucin gene expression.

**Fig 3.**
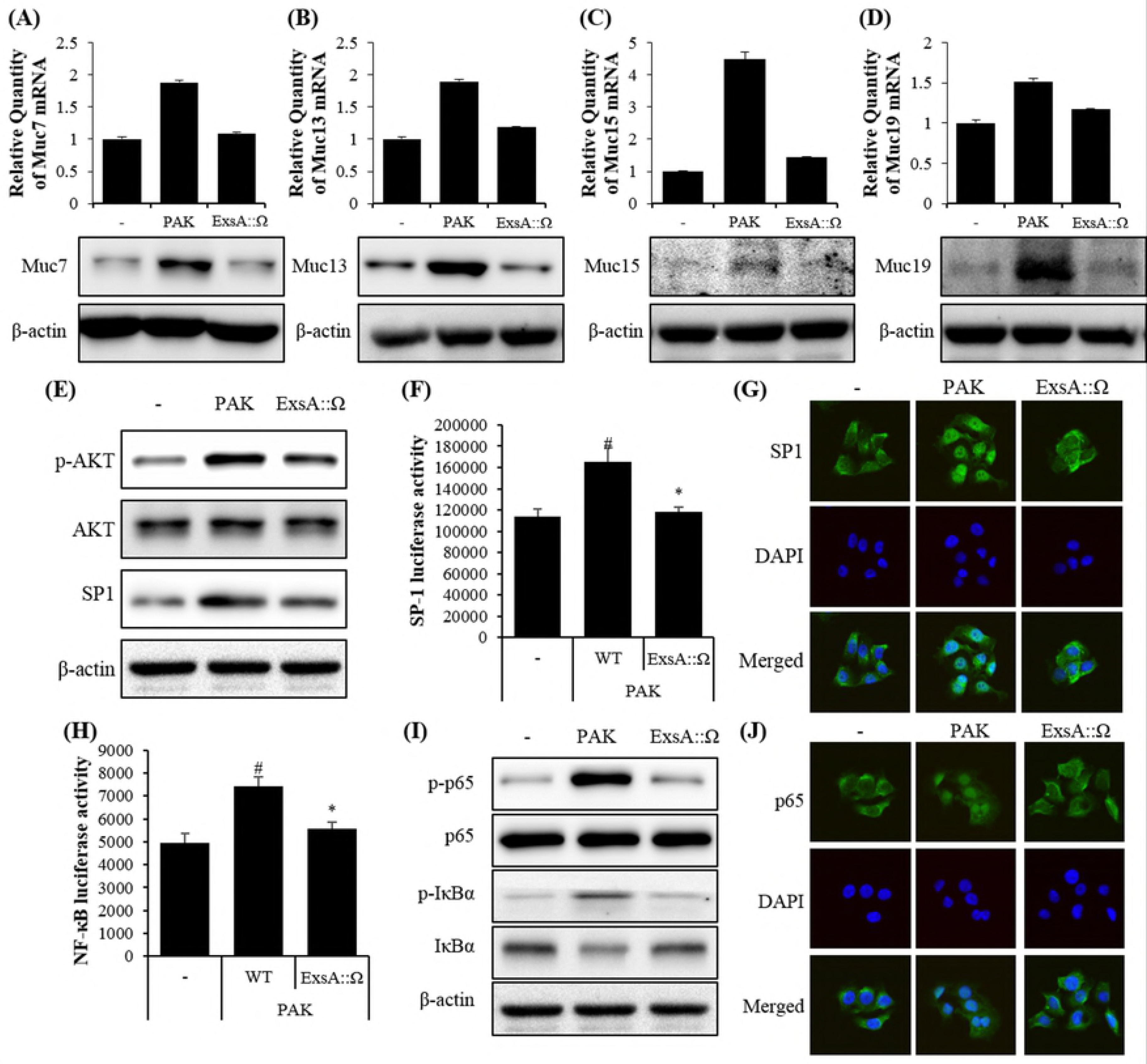
ExsA::Ω affects Muc7, Muc13, Muc15, and Muc19 expression in A549 and H292 cells. The expression levels of Muc7 (A), Muc13 (B), Muc15 (C), and Muc19 (D) mRNA and protein were determined by qRT-PCR and western blotting, respectively. A549 cells were treated with PAK and ExsA::Ω for 1 h, followed by 2 washes for 15 min and further treatment for 4 h. PAK exposure increases the phosphorylation of AKT (E, F) and p65 (H, I) compared with the vehicle control group in A549 cells. Transcription of SP1 (G) and NF-kB (p65) (J) into the nucleus were confirmed by immunocytochemistry (ICC) in H292 cells. The data are presented as the mean ± SD. “–”, A549 cells treated with PBS; PAK, A549 cells infected with *P. aeruginosa*; ExsA::Ω, A549 cells infected with ExsA::Ω (no T3SS). #Significantly different from the non-infected control cells, \**P* < 0.05, significantly different from PAK-only-infected cells.

**Fig 4.**
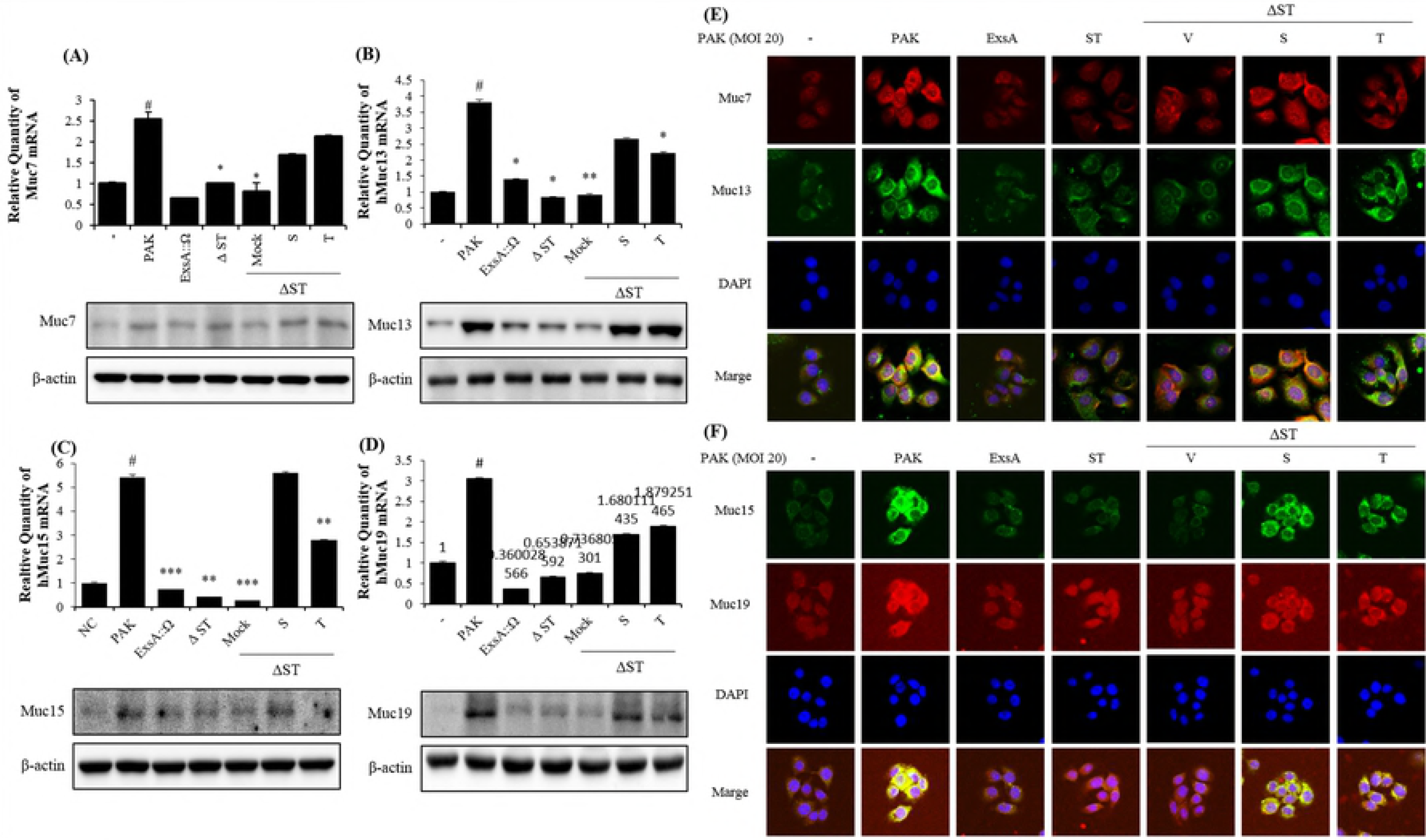
ExoS and ExoT are required and each sufficient to induce Muc7, Muc13, Muc15, and Muc19 expression in A549 and H292 cells. (A–D) Muc7, Muc13, Muc15, and Muc19 mRNA and protein expression as determined by qRT-PCR and western blotting, respectively. (E–F) The cellular localization of Muc7, Muc13, Muc15, and Muc19 as determined by immunocytochemistry in H292 cells. “–”, control A549 and H292 cells treated with PBS only; PAK, A549 or H292 cells infected with *P. aeruginosa*; ExsA::Ω, A549 or H292 cells infected ExsA::Ω (no T3SS); ΔST, PAK-ΔST mutant; Mock, PAKΔSTmt-pUCP18; S, PAKΔSTmt-pUCP18-PAKexoS; T, PAKΔSTmt-pUCP18-PAKexoT (MOI = 200) #Significantly different from the non-infected control cells, \**P* < 0.05, \*\**P* < 0.01, \*\*\**P* < 0.001, significantly different from PAK-only-infected cells.

### ExoS and ExoT of T3SS are required for the increase in Muc7, Muc13, Muc15, and Muc19 and inflammatory cytokine expression in A549 cells

Next, we assessed the roles of ExoS and ExoT in the induction of mucin expression by using a mutant strain defective in these two proteins (ΔST) and complementation strains of ΔST with either of the proteins restored (S, expressing ExoS only, and T, expressing ExoT only). We observed that ExoS and ExoT are critical for the expression of the 4 mucins gene in A549 cells as ΔST could not induce Muc7, Muc13, Muc15, and Muc19 expression, while the induction of mucin expression was partially restored in the complementation strains restored their expression induction. Real-time PCR and ELISA indicated that IL-6 expression was restored to approximately 90% and 80% of the PAK-induced level in case of infection with ExoS and ExoT, respectively, and TNF-α was restored to about 60% and 55% of the PAK-induced level in case of infection with ExoS and ExoT, respectively (S3A–D Fig). Thus, in accordance with previous reports, the expression of Muc7, Muc13, Muc15, and Muc19 and proinflammatory cytokines induced by PAK in A549 cells, requires ExoS and ExoT. In cells treated with ExsA::Ω, the increase in IL-6 and TNF-α was suppressed as compared to cells treated with wild-type PAK. ΔST totally lost the ability to induce IL-6 and TNF-α, while this ability was restored in the complementation strains expressing either ExoS (S) or ExoT (T) (S3A–D Fig).

### ExoS and ExoT of T3SS are required for the increase in Muc7, Muc13, Muc15, and Muc19 expression via NF-κB p65 and AKT phosphorylation in A549 cells

Next, we investigated whether ExoS and ExoT are related to known SP1/AKT and NF-κB signals, which are involved in pneumonia and lung diseases, using the mutant strains. The results showed that S and T induced the activation of the SP1/AKT pathway and AKT and p65 phosphorylation in A549 cells nearly to the level the wild-type PAK strain did, while ΔST completely the ability to do so (Fig 5A–D). In particular, it was confirmed that increased SP1, known as a transcription factor for AKT and increases the phosphorylation of IκBα (Fig. 5A–D). These results indicated that ExoS and ExoT are important virulence factors of *P. aeruginosa* and are very important for the induction of the inflammatory response in host cells.

**Fig 5.**
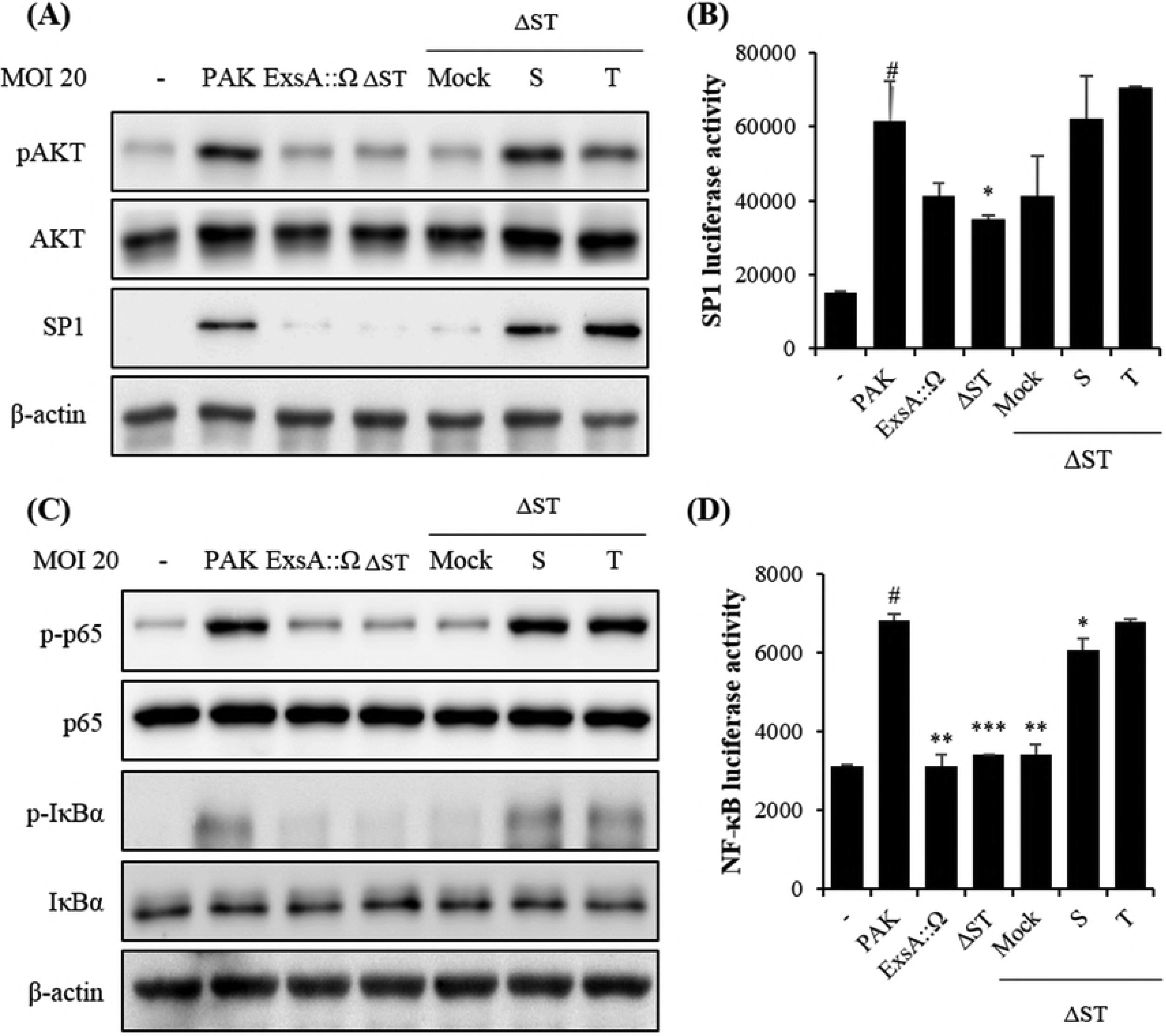
ExoS and ExoT are required and each sufficient to increase NF-κB and AKT signaling in A549 cells. A549 cells were infected with the indicated strains for 1 h. (A) AKT and p-AKT expression as detected by western blotting. Quantification of p-AKT and SP1 expression using RAS-4000. (B) A549 cells were transfected with SP1 luciferase (Luc) reporter plasmid (0.1 μg). (C) NF-κB expression as detected by western blotting. Quantification of p-p65 and p-lκB-α expression using RAS-4000. β-actin was used as the internal control. (D) A549 cells were transfected with expression NF-κkB luciferase (Luc) reporter plasmid (0.1 μg). At 24 h after transfection, A549 cells were treated with the strains at MOI = 20 for 1 h, and then, luciferase activity was measured. The data were normalized to β-galactosidase activity. All data are representative of 3 independent experiments. Luciferase activities were measured 24 h after the transfection. “–”, control A549 cells treated with PBS only; PAK, A549 cells infected with *P. aeruginosa;* ExsA::Ω, A549 cells infected with ExsA::Ω (no T3SS); ΔST, PAK-ΔST mutant; Mock, PAKΔSTmt-pUCP18; S, PAKΔSTmt-pUCP18PAKexoS; T, PAKΔSTmt-pUCP18PAKexoT (MOI = 20).

### Effect of PAK infection in a pneumonia mouse model and the roles of ExoS and ExoT therein

We determined the PAK infection level in a C57BL/6 mouse model. It is known that PAK infection at specific MOI induces an inflammatory in mice. Accordingly, we tested PAK infection at different MOIs to select the most appropriate one. Pure LPS was used as a control, and mice were subjected to infection with PAK alone (PAK) or together with LPS treatment (PAK+LPS). PAK or LPS was diluted with PBS to inject into mice. Experiments were conducted using 7-8 mice per group. The protein levels of Muc13, Muc15, and Muc19 were increased in lung tissues of PAK+LPS-infected mice (S4A–C Fig). Hematoxylin and eosin (H&E) staining of lung tissues revealed that inflammatory cells were increased in PAK-infected and in PAK+LPS-treated mice. In particular, treatment of PAK+LPS together with LPS and PAK alone resulted in a much greater increase in inflammatory cells. (S5 Fig).

When we tested the ΔST, S, and T strains, Muc7, Muc13, Muc15, and Muc19 mRNA and protein expression were significantly increased by S and T, but not ΔST, as indicated by qPCR and western blotting (Fig 6A–E). Similar findings were achieved for proinflammatory cytokine expression (data not shown). Thus, mucin gene expression in the mouse lungs in response to *P. aeruginosa* infection is controlled by ExoS and ExoT.

**Fig 6.**
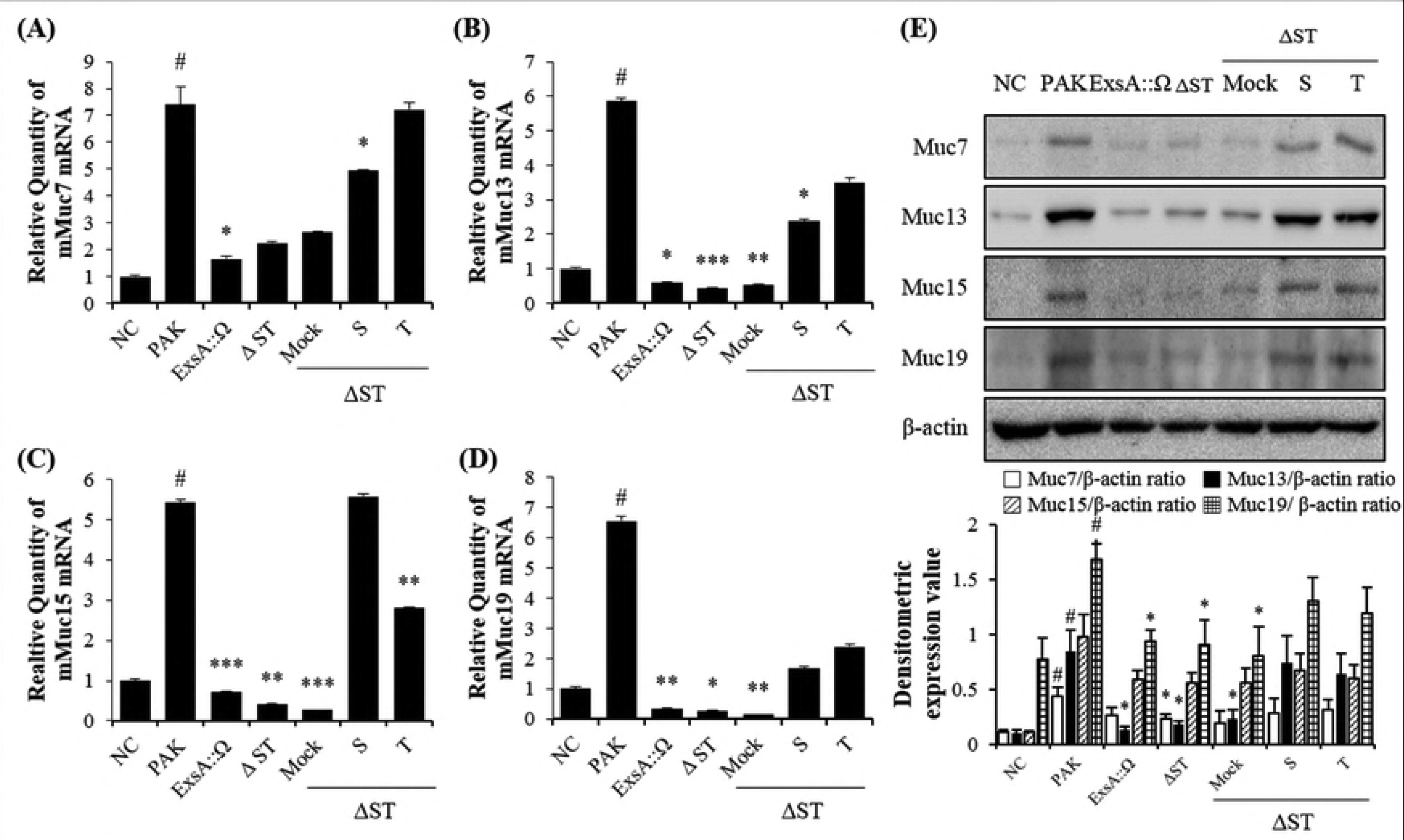
ExoS and ExoT are required to induce Muc7, Muc13, Muc15, and Muc19 expression in lung tissue. (A–D) Muc7, Muc13, Muc15, and Muc19 mRNA levels as determined by qRT-PCR. (E) Western blot showing Muc7, Muc13, Muc15, and Muc19 protein expression and quantitative data. NC, control mice treated with PBS only; PAK, mice infected with *P. aeruginosa;* ExsA::Ω, mice cells infected ExsA::Ω (no T3SS); ΔST, PAK-ΔST mutant; Mock, PAKΔSTmt-pU CP18; S, PAKΔSTmt-pU CP18PAKexoS; T, PAKΔSTmt-pUCP18PAKexoT (MOI 2.5 × 10^6^) #Significantly different from the non-infected control cells, \**P* < 0.05, \*\**P* < 0.01, \*\*\**P* < 0.001, significantly different from PAK-only-infected cells.

### Roles of ExoS and ExoT in AKT activation in mice

Proinflammatory cytokine production was increased through the T3SS of PAK, and this involves AKT activation in pneumonia mouse models. To confirm the role of SP1 in S- and T-infected pneumonia model mice, we evaluated the level of AKT phosphorylation by western blot analysis using antibodies to SP1, AKT, and pAKT. Non-treated as well as ΔST-treated cells displayed weak AKT phosphorylation, whereas S- and T-treated monolayers showed a significant increase in SP1 translocation, which was detectable as of 30 min and was sustained for 1 h (Fig 7A).

**Fig 7.**
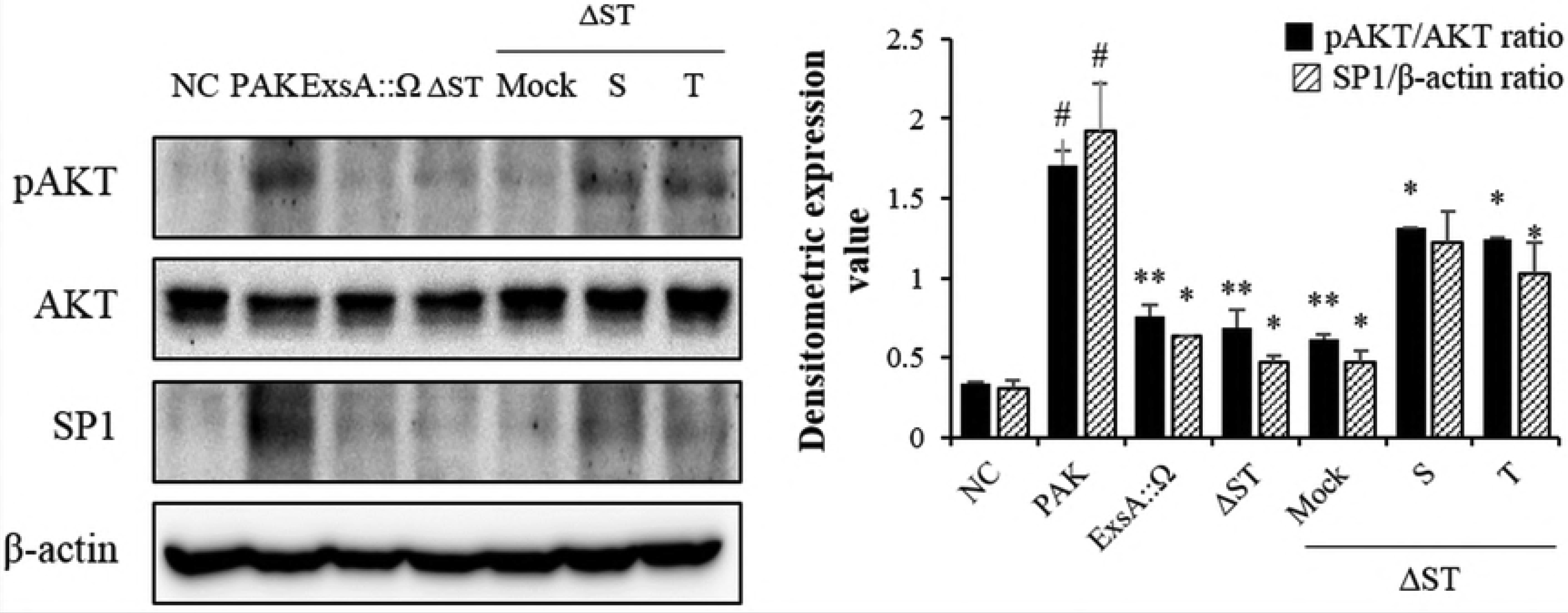
ExoS and ExoT are required and each sufficient to increase AKT/SP1 signaling in lung tissue. Western blot showing pAKT, AKT, and SP1 protein expression and quantitative data. NC, control mice treated with PBS only; PAK; ExsA::Ω (no T3SS), ΔST, S, and T (MOI 2.5 × 10^6^). #Significantly different from the non-infected control cells, \**P* < 0.05, \*\**P* < 0.01, significantly different from PAK-only-infected cells.

### IKK-α and IKK-β are necessary for NF-κB phosphorylation in response to S and T infection

The IκB kinase enzyme complex is part of the upstream NF-κB signal transduction cascade. The IκB kinase (IKK) is an enzyme complex that is involved in propagating the cellular response to inflammation [25]. Small interfering (si)RNA is one of the experimental tools for functional analyses. We used siRNAs to knock out IKKα and IKKβ expression. The presence of control siRNA (siNC) did not affect the increase in muc7, Muc13, Muc15, and Muc19 expression induced in A549 cells by PAK infection. However, knockout of IKKα and IKKβ suppressed the increases in muc7, Muc13, Muc15, and Muc19 by 50% (S6A–D Fig). ELISA of proinflammatory cytokines IL-6 and IL-8 revealed similar effects as those observed for the mucins (S7A, B Fig).

Next, we conducted experiments using the strains S and T. A549 cells infected with PAK, ExoS, or ExoT after siNC transfection induced increased muc7, Muc13, Muc15, and Muc19 expression as compared to the negative control. However, in A549 cells infected with PAK, ExoS, or ExoT after knockout of IKKα and IKKβ, the increases in Muc7, Muc13, Muc15, and Muc19 were suppressed by 40–60% (Fig 8A–D).

**Fig 8.**
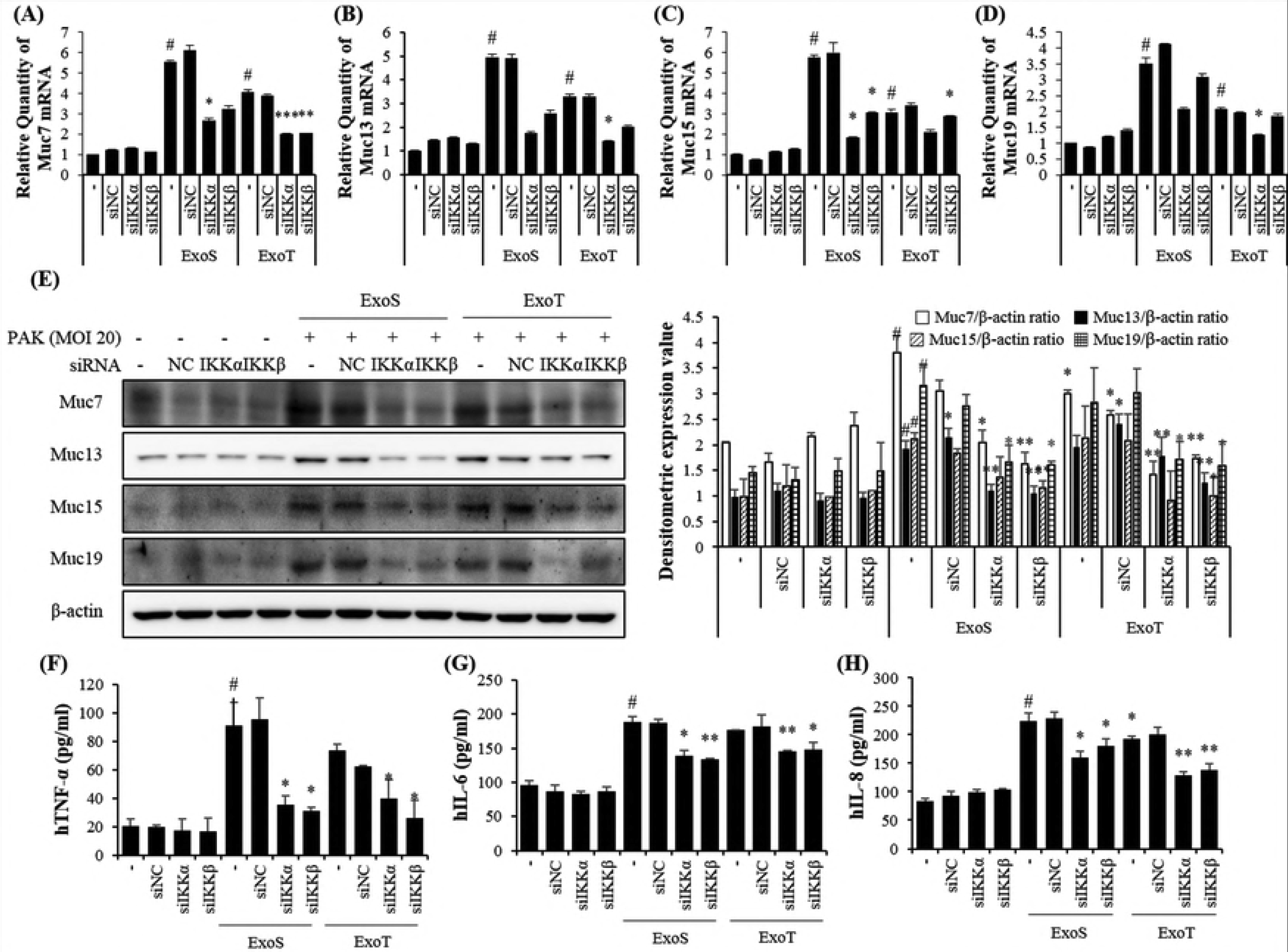
Muc7, Muc13, Muc15, Muc19, and proinflammatory cytokine expression require the NF-κB pathway activated via IKK-α and IKK-β in A549 cells. (A–D) qRT-PCR analyses showing Muc7, Muc13, Muc15, and Muc19 mRNA expression upon the addition of siRNAs targeting IKK-α and IKK-β. (E) Western blot showing Muc7, Muc13, Muc15, and Muc19 protein expression upon the addition of siRNAs targeting IKK-α and IKK-β as determined by western blotting and quantitative data. (F–H) Cytokine levels as determined by ELISA. Data are the mean ± SEM of 3 different experiments performed in duplicate.

## Discussion

T3SS gene expression is induced by contact with eukaryotic cells or under specific environmental conditions [26]. ExoS and ExoT are toxins with adenosine diphosphate ribosyltransferase and Rho guanosine triphosphatase activities [27]. They are similar in structure, but the toxicity of ExoT is less potent than that of ExoS [28,29]. When activated, these proteins destroy the actin filaments that make up the cytoskeleton of the host, thus inhibiting phagocytosis and finally killing the host cell [30]. We found that S and T complementation strains were extremely efficient in lysing eukaryotic cells, and this observation corresponded with their highly toxic phenotypes in a pneumonia mouse model. S and T were found to be as damaging as the T3SS-proficient PAK strain in mice. Histological analysis *in vivo* showed that S- and T-infected lungs had damage lesions.

*P. aeruginosa* ExoS and ExoT were suggested to be associated with lung injury and mucus accumulation in mice [31,32], but these types of virulence mechanisms have never been investigated at the molecular level. The way in which these major virulence determinants of *P. aeruginosa* function to cause severe disease during infection was not clear until date. Therefore, the effects of NF-κB and SP1/AKT activities on the expression of Muc7, Muc13, Muc15, and Muc19 in S- and T-infected pneumonia model mice were investigated. Expression and delivery of Muc7, Muc13, Muc15, and Muc19 were found to be improved by S and T infection. Examination of infected A549 cells revealed that the expression of Muc7, Muc13, Muc15, and Muc19 is dependent on NF-κBp65 and SP1/AKT phosphorylation. In particular, overexpression of the inflammatory cytokines (IL-6, IL-1β, and TNF-α) in response to activation of ExoS- and ExoT-induced NF-κB and AKT was associated with the pathogenesis mechanism in the pneumonia mouse model. In addition, proinflammatory cytokine expression as well as expression of Muc7, Muc13, Muc15, and Muc19 was increased upon infection with strains S and T. Our results suggest that ExoS and ExoT play important roles in the genetic etiology associated with acute and chronic infections.

*P. aeruginosa* is known to have an important influence on the activation of NF-κB and SP1 in pulmonary diseases and to regulate the production of cytokines, matrix metalloproteinases (MMPs), and mucins [4,33]. Muc7, Muc13, Muc15, and Muc19 gene and protein expression is facilitated by several cytokines, including TNF-α and IL-1β, secreted by macrophages [8,34]. In previous reported that IL-6, IL-8, and TNF-α secreted by lung lymphocytes specifically induce the secretion of Muc7, Muc13, Muc15, and Muc19 to promote destruction of lung tissue [35,36]. Muc7, Muc13, Muc15, and Muc19, IL-1β, and TNF-α expression levels in response to infection were significantly reduced in A549 cells transformed with siRNA targeting IKK-α and IKK-β. This shows that Muc7, Muc13, Muc15, and Muc19 are required for the expression of proinflammatory cytokines. Since mucin expression is known to increase respiratory irritation in pulmonary infections, it further implies cytokine-induced pathogenicity, as observed in the lung biopsy specimens.

The results generated in the current study increase our understanding of the pathogenesis of pneumonia by demonstrating that the T3SS, and specifically, ExoS and ExoT, disrupt the host response in the lungs. Most importantly, ExoS and ExoT of T3SS induce the activation of NF-κB and AKT, and are able to induce the expression of Muc7, Muc13, Muc15, and Muc19 in the lungs. Although these 4 mucins have not been studied thoroughly yet, they are considered important inflammatory mediators in inflammation of the respiratory tract. Treatment of chronic respiratory inflammation is limited to vaccination against commonly occurring respiratory pathogens, pharmacological bronchodilation, or respiratory infection antibiotics to alleviate dyspnea. Our research results provide a basis for future therapies to prevent and confound lung immunopathology through increasing our understanding of the molecular mechanism of pneumonia.

## Acknowledgments

This study was supported by the National Research Foundation of Korea (NRF) grant funded by the Ministry of Science, ICT & Future Planning (NRF-2017R1A2B2011555) and awarded to the Korea Research Institute of Bioscience and Biotechnology Research Initiative Program (KGM 1221814) of the Republic of Korea.

## Supporting information

**S1 Fig.**
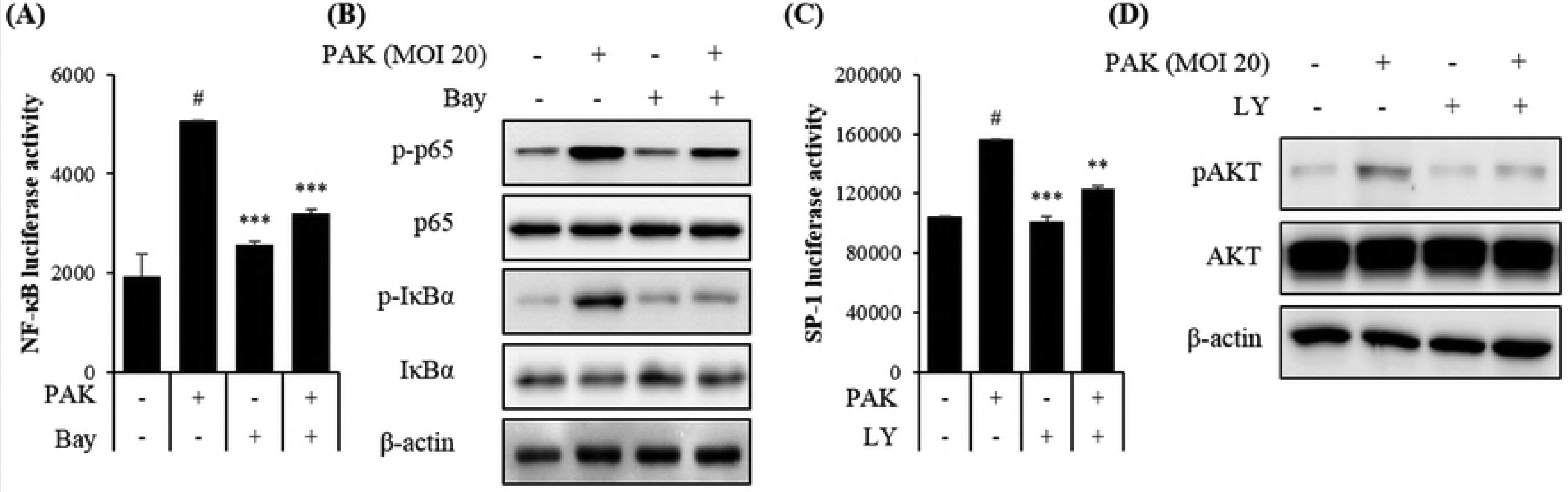
Effects of PAK exposure on the phosphorylation of NF-κB and SP1/AKT in A549 cells.

**S2 Fig.**
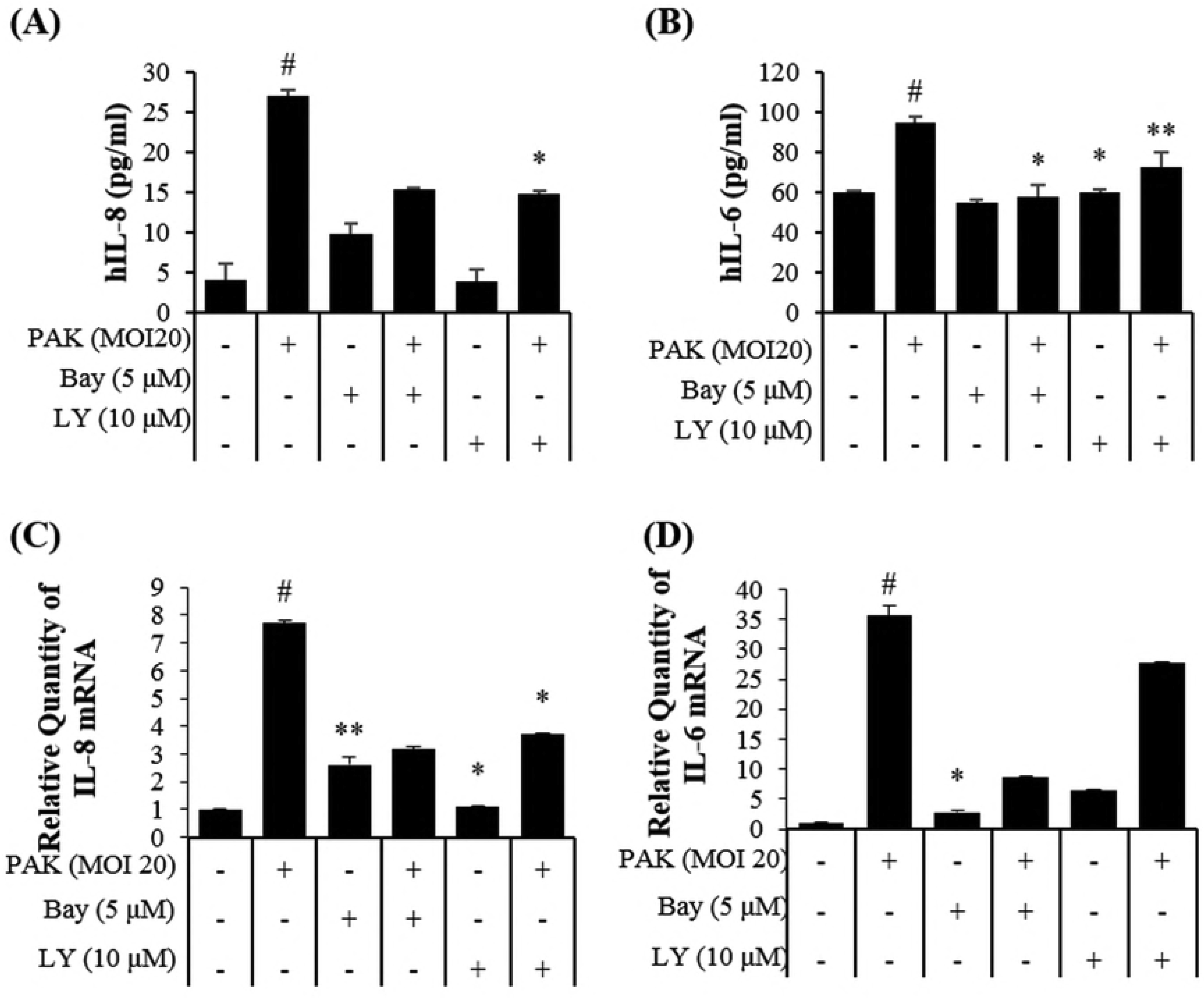
Effects of inhibitors on the expression of proinflammatory cytokines in PAK infected A549 cells.

**S3 Fig.**
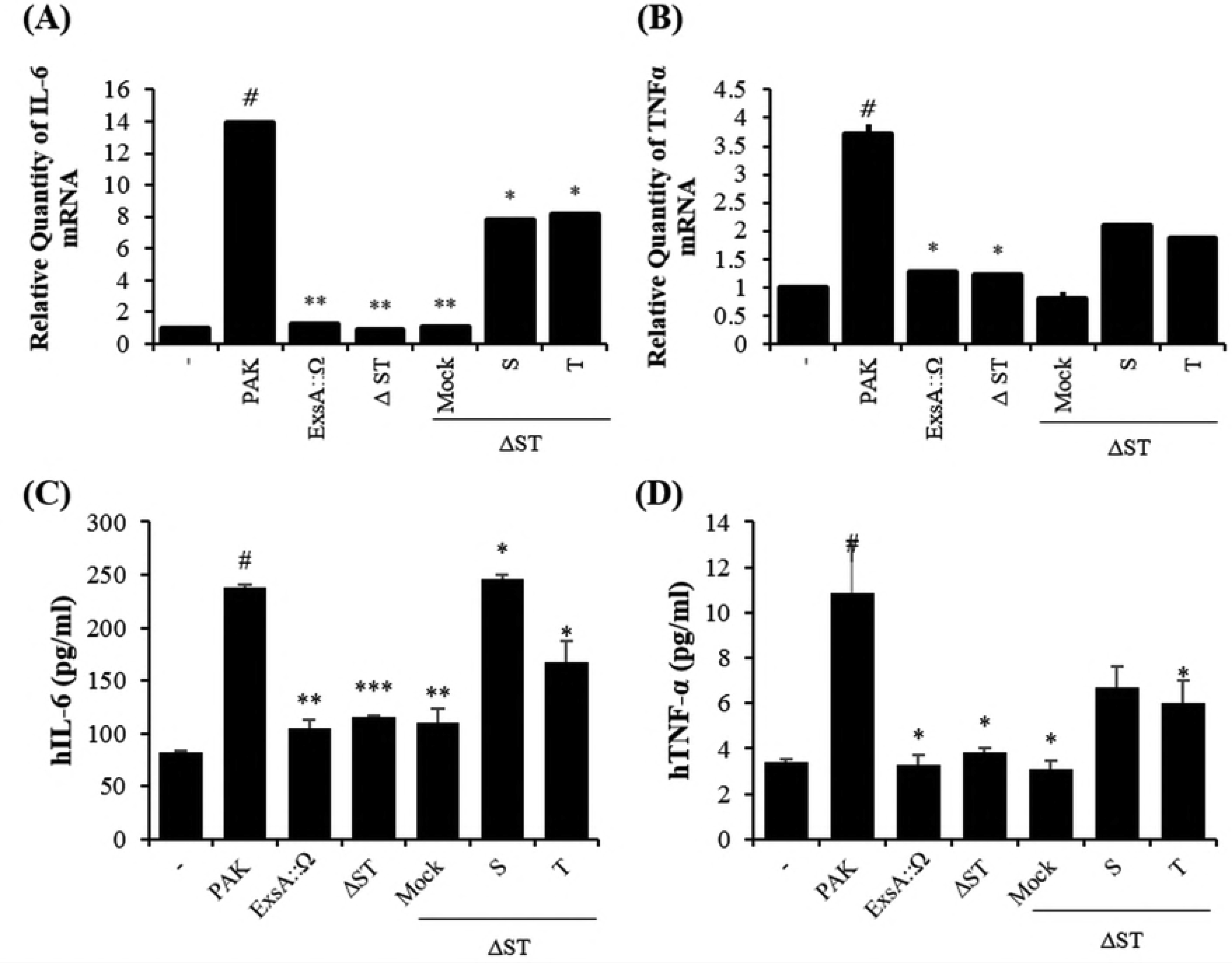
Effects of ExoS and ExoT exposure of proinflammatory cytokines in PAK infected A549 cells.

**S4 Fig.**
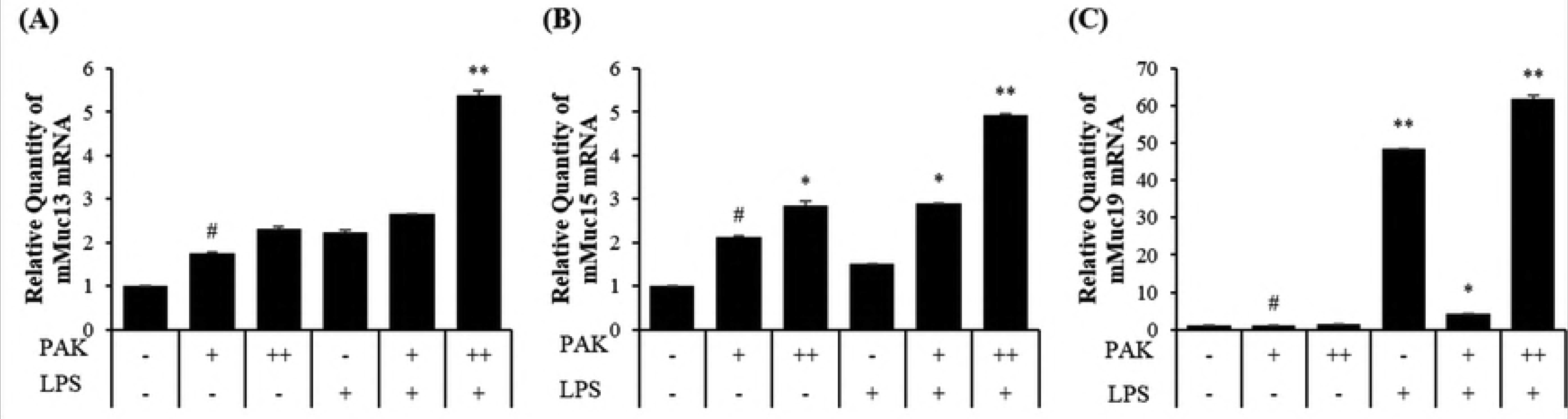
Effects of PAK and LPS exposure of the Muc13, Muc15, and Muc19 in mice.

**S5 Fig.**
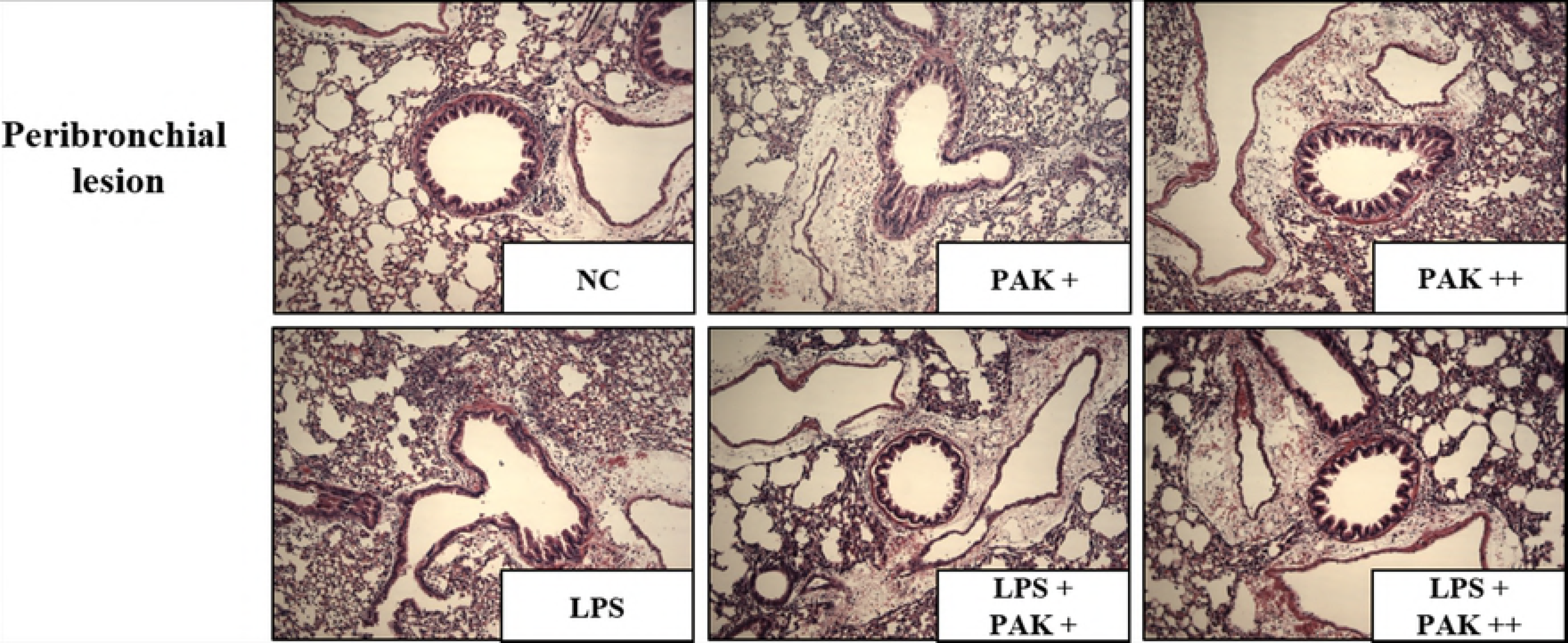
Histopathology of mouse lungs 20 h after inoculation with *P. aeruginosa* and LPS.

**S6 Fig.**
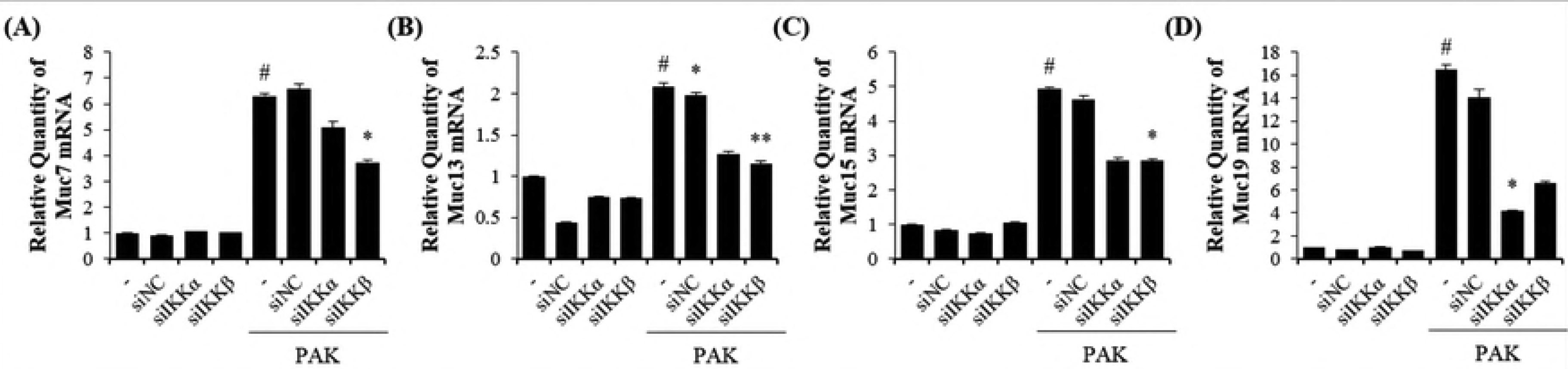
Muc7, Muc13, Muc15, and Muc19 in A549 after siRNA-mediated downregulation of IKK-α and IKK-β.

**S7 Fig.**
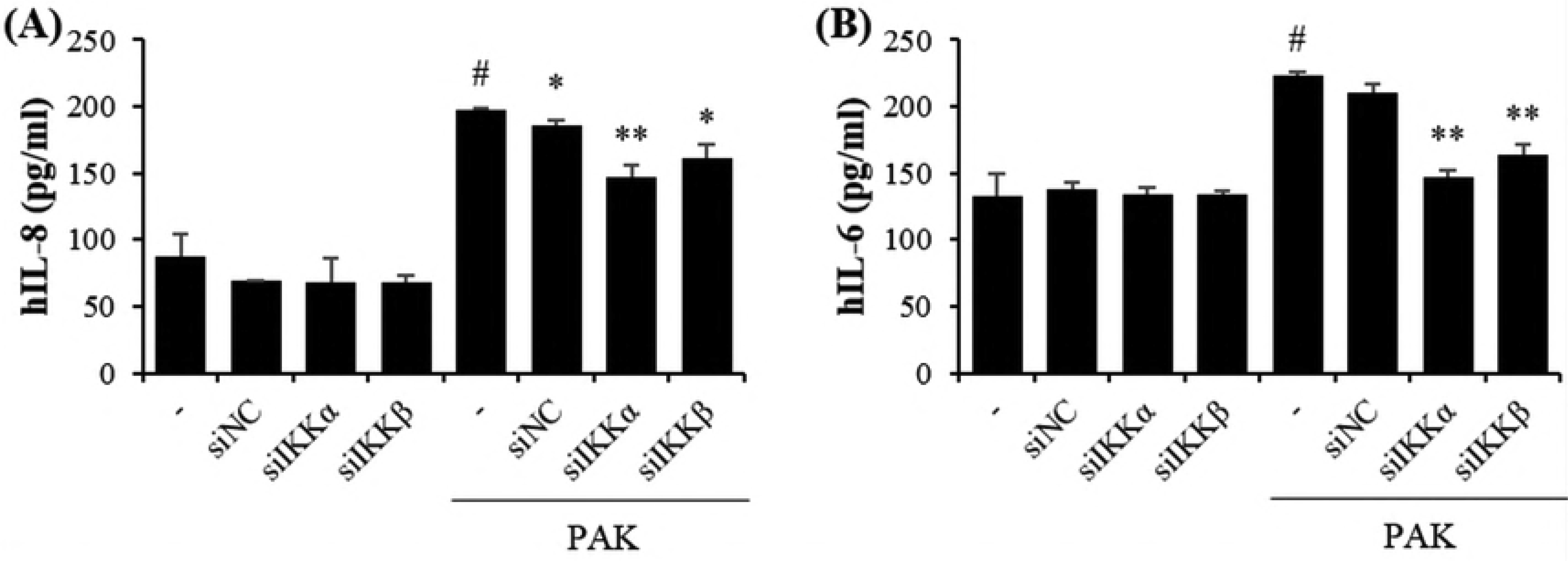
Proinflammatory cytokine in A549 after siRNA-mediated downregulation of IKK-α and IKK-β.

